# DNA extraction free whole genome sequencing of bacteriophage genomes from a single plaque

**DOI:** 10.1101/2025.10.21.683754

**Authors:** Brenna Fox, Jasmin Chahal, Patrick Lypaczewski

**Affiliations:** Department of Microbiology and Immunology, McGill University, Montreal, Canada

## Abstract

DNA sequencing is at the core of genome characterization, proteomics, and identification of novel organisms. For microorganisms such as bacteriophages, sequencing their DNA can provide key insights into their tropism, infectivity, and virulence. There remains however a critical lack of rapid sequencing techniques with the traditional process of replating and incubating individual plaques, collecting lysate, extracting DNA, preparing the DNA library, and sequencing that is labor intensive. Herein, we demonstrate the use of an adapted Nanopore Rapid PCR Barcoding protocol to sequence the bacteriophage genome directly from individual plaques. This technique provides sequencing genome assemblies with 99.88-100% (mean 99.97%) average nucleotide identity (ANI) scores when compared to the traditional methods involving phage amplification, extraction, and sequencing using Illumina. The optimization of bacteriophage identification by the technique of *tagmentation* directly to isolated plaques will enable rapid and cost-effective sequencing of novel phages.

## Introduction

Bacteriophages (phages) are the most abundant organisms on the planet and have been gaining traction as powerful microorganisms that can be used in biotechnology (Mushegian, 2020). One promising application of phages is their potential to combat antimicrobial resistance (AMR). AMR poses a significant threat to human health by rendering antibiotics ineffective against bacterial infections (Salam et al., 2023). As phages have specific tropism for bacteria and can selectively kill them using a lytic life cycle, phages have become promising as antimicrobial treatments in both medicine and agriculture (Elfadadny et al., 2024; Zhang et al., 2022) or as potential modulators of the microbiome (Pottie et al., 2024).

Due to phage specificity, large library collections are required to screen for potential therapeutic use. In an effort to increase phage discovery, programs such as The Science Education Alliance-Phage Hunters Advancing Genomics and Evolutionary Science (SEA-PHAGES) crowd-source the discovery and characterization of novel phages while also promoting student involvement in science (Heller et al., 2024). Through the participation of undergraduate students enrolled in universities worldwide, over 28,000 novel phages have been identified. However, only around 5,000 of these phages have been sequenced (Gauthier & Hatfull, 2024). Due to current limitations of sequencing costs and time, only a fraction of isolated phages is sequenced each year as is the case in the SEA-PHAGES program. The complexities involved in amplifying, extracting and purifying DNA from phages lead to a bottleneck of unexplored, potentially valuable phages in all areas of phage research, including but not limited to the SEA-PHAGES program. This bottleneck leads to a detachment of the personalized learning approach that SEA-PHAGES aims to implement, as students rarely sequence the phages that they isolate.

The current standard for DNA sequencing of phages requires the isolation of single plaques, followed by overnight or multi-hour amplification in either liquid or solid culture using a suitable host, followed by the filtration of phage particles prior to DNA extraction. This purified DNA can then be sequenced on different platforms including Illumina, PacBio or Oxford Nanopore Technologies (ONT). Due to its inherent high accuracy, Illumina is still preferred by multiple groups, including the SEA-PHAGES programs (Rihtman et al., 2016). However, the use of ONT sequencing has been investigated by several groups and the accuracy of ONT sequencing is now approaching Illumina’s (González-Escalona et al., 2019). Additionally, the implementation of Illumina sequencing is generally more costly compared to ONT sequencing. Further benefits of ONT sequencing include the ability to sequence long reads spanning multiple kilobases, while traditional methods of DNA sequencing are only capable of sequencing short fragments of DNA in the range of 50-500 bases. This limitation can result in difficulties with sequencing and assembling DNA with repetitive regions, as it can lead to assembly gaps. Since ONT can read the entirety of repetitive regions, this technology can sequence DNA without the risk of assembly gaps. In addition, ONT DNA sequencing can be accomplished in-house with low initial costs and portable instruments (Zhang et al., 2024; Zheng et al., 2023). Due to the efficiency in time and cost of ONT sequencing, this technology is ideal for use in student labs, in the field or in resource-limited settings.

A common approach to sequencing using ONT is to prepare DNA for sequencing using the *Rapid* family of library preparation kits. In kits such as the *Rapid Barcoding Kit V14* (SQK-RBK114-24), purified DNA is fragmented and tagged simultaneously using transposases (tagmentation) that contain distinct barcodes flanked by proprietary click-chemistry reactive molecules. After barcoding, *Rapid Adaptors* (RA) are added to the samples. These adaptors are oligonucleotides protein complexes required for pore docking and sequencing. They contain a complementary click-chemistry molecule allowing for the attachment to the target DNA (Oxford Nanopore, 2024). Barcoding of samples is advantageous when multiple samples are sequenced on the same run using ONT such as for example DNA from phage samples isolated from different locations.

While these transposase-based preparation kits are intended to reduce handling-time compared to traditional sequencing kits, they require purified extracted DNA as input material. Herein, we investigate the possibility of adapting the rapid sequencing kits for use with crude phage material originating directly from single plaque picks. We describe the optimization of the protocol and demonstrate the successful whole genome sequencing of multiple phages isolated directly from single plaque picks, bypassing phage amplification or DNA extraction steps.

## Methods

### Bacterial Culture

Host bacteria *Arthrobacter globiformis* B-2979 was cultured by inoculating 10mL PYCa (Peptone, Yeast, Calcium agar) media. Incubation at 30°C for 48 hours resulted in a turbid culture, indicative of bacterial growth. A negative control of PYCa media inoculated with no bacteria was also performed.

### Plaque Assay Plates

Individual phage plaques were obtained by performing plaque assays with the host bacteria *A. globiformis* (Supplementary Figure S1). Briefly, 100-200µL of *A. globiformis* with 10µL of serially diluted phage stock in phage buffer (10mM Tris pH 7.5, 10mM MgSO4, 68mM NaCl, 1mM CaCl2) was incubated for 10 minutes to allow for attachment, and then subsequently suspended in PYCa liquid top agar and plated. In total, 21 different bacteriophages were plated and used for this experiment (Supplementary Table S1). These plates were incubated with inversion at 30°C for 24 hours.

### Bacteriophage Direct Isolation and Amplification Using Non-Barcoding Primers

Zones of clearance (plaques) from bacteriophage plaque assays were picked in a sterile environment by stabbing the top layer of agar with a 20μL Axygen filtered pipette tip and resuspended in 50μL TE (10 mM tris pH 8, 0.1mM EDTA). After brief vortexing, 3μL of each TE sample was transferred to a PCR tube along with 1μL of transposase (FRM from Nanopore SQK-RPB114.24 or RB#1-24 from Nanopore SQK-RBK114.24) to allow for fragmentation (Figure 1A). This MuA transposase inserts a known MuA recognition sequence (Green), a barcode sequence (Red), and a barcode flanking region (Oxford Nanopore, 2024; Yanagihara & Mizuuchi, 2003). A positive control consisting of 5ng of purified phage DNA and a negative control consisting of sterile TE were each combined with 1μL of transposase. The samples were incubated in a thermocycler at 30°C for 2 minutes followed by 80°C for 2 minutes.

**Figure 1.**
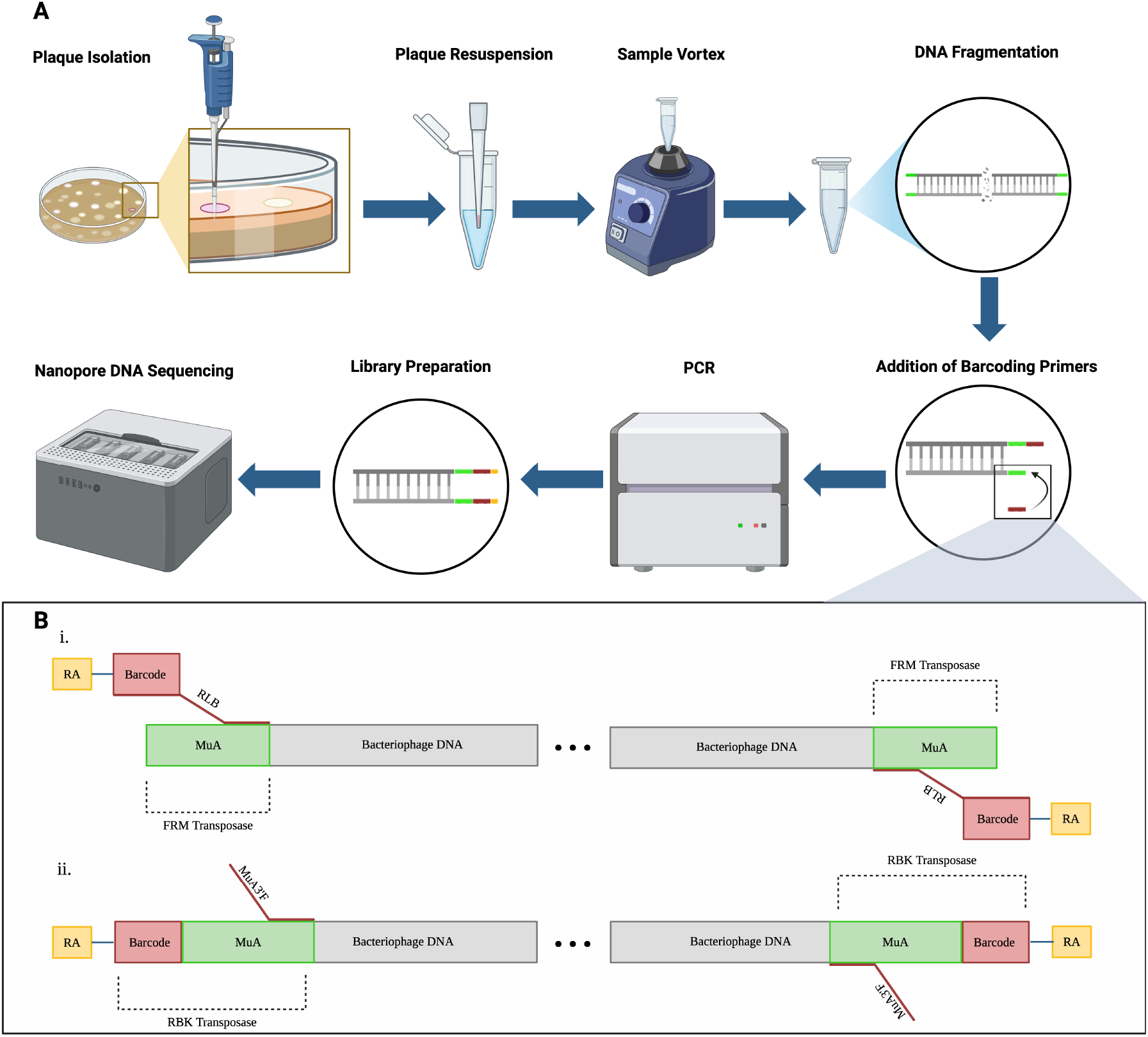
Protocol flowchart and illustration of DNA binding: Nanopore Rapid PCR Barcoding Kit and prescreen reagents. A) Flowchart of the direct-from-plaque DNA sequencing protocol. Individual plaques are picked from the top agar layer using a pipette tip and resuspended in TE buffer. Following brief vortexing, the plaque suspension undergoes transposase-mediated fragmentation (tagmentation), where the transposase enzyme simultaneously cuts the DNA and inserts its pre-loaded cargo into the target DNA. The fragmented DNA is then amplified by PCR using primers that bind to the transposase-inserted sequences. After amplification, rapid adaptors (RA) are attached via click chemistry to the complementary moiety on the RLB primers (yellow), and the library is sequenced on an ONT platform. B) Schematic representation of transposase-inserted DNA sequences and primer binding sites. Two transposase enzymes are shown: FRM (from the Rapid PCR Barcoding Kit, top) and RBK (from the Rapid Barcoding Kit, bottom). Both transposases insert a common MuA recognition sequence (green) into the phage DNA during fragmentation. The RBK transposase cargo additionally contains a barcode sequence (red) and RA complementary moiety (yellow) for attachment of the fragmented DNA to sequencing adaptors. The FRM transposase does not include a barcode. (i) The RLB primer (from ONT’s Rapid PCR Barcoding Kit) binds to the MuA recognition sequence inserted by either FRM or RBK transposase. Unlike MuA3’F, the RLB primer contains an integrated barcode sequence and proprietary click chemistry moieties that enable rapid attachment to the Rapid Adaptor (RA). (ii) The MuA3’F primer (an in-house diagnostic primer) binds to the MuA recognition sequence. This primer does not contain barcode sequences or click chemistry groups and is used for pre-screening to confirm successful tagmentation before committing to more expensive sequencing primers. Created in BioRender. Fox, B. (2025) https://BioRender.com/7ag1ntl

Subsequently, 1μL of each fragmented mix was transferred to a clean PCR tube along with 10μL SuperFi II DNA Polymerase Master Mix (ThermoFisher Scientific), 8.8μL nuclease-free water, and 1.0μM of a pre-screen diagnostic primer (MuA3’F: 5’-CGT TTT TCG TGC GCC GCT TC -3’). This primer is designed to bind the MuA 3’ region, a known sequence inserted by the transposase (Figure 1B). The same primer serves as forward and reverse primer and amplifies DNA fragments tagged on both 5’ and 3’ by the transposase. PCR cycling was performed according to the manufacturer’s instructions for the SuperFi II DNA Polymerase Protocol (ThermoFisher Scientific) allowing a fragmentation size reaching up to 15kb (5-6 minutes extension time) with an additional 5-minute final extension. To confirm the amplification of samples, 5μL of each sample were loaded on a 1-1.5% agarose gel in addition to the remaining fragmented unamplified DNA and imaged after 50 minutes at 90V.

### Bacteriophage Direct Isolation and Sequencing Using ONT Barcodes

Following successful optimization using MuA3’F pre-screen primers, phage plaques were picked in a sterile environment by stabbing the top layer of agar with an Axygen 20μL filtered pipette tip and resuspended in 50μL of TE (10 mM tris pH 8, 0.1mM EDTA). The samples were vortexed briefly followed by transposase mediated fragmentation and PCR amplification as described above, replacing the non-barcoding MuA3’F primer with 1.0μL RLB #1-24 sequencing primer (SQK-RPB114.24). Additionally, 8.0μL nuclease-free water was added to the PCR reaction as opposed to 8.8μL, allowing for the total volume of 20μL to remain constant.

Post-PCR, 10μL of each sample was pooled into a 1.5mL Eppendorf tube with a 0.8X volume ratio of Ampure XP beads (Beckmann). Beads were incubated at room temperature for 10mins and pelleted on a magnetic rack followed by two 1mL 80% ethanol washes. Residual ethanol was removed, and the DNA was eluted in 25μL Elution Buffer (SQK-RPB114.24) for 5 minutes at room temperature. To remove residual green loading dye, two rounds of cleanup were performed on reactions that used the Platinum SuperFi II Green PCR Master Mix.

The purified PCR pool was used to generate a sequencing library by incubating the pooled DNA with Rapid Adaptor (RA, SQK-RPB114.24). Sequencing was performed on a previously used and washed PromethION R10.4.1M flow cell nearing end-of-life as defined by the manufacturer (100-1000 pores) to minimize costs, functionally similar to sequencing on new MinION or Flongle flow cells.

### Phage DNA Extraction from Lysates and Whole Genome Sequencing

To create a reference genome for previously non-sequenced phages, highly concentrated plaque assays were produced. The plates were flooded in 8mL of phage buffer (10mM Tris pH 7.5, 10mM MgSO4, 68mM NaCl, 1mM CaCl2) incubated for 4 hours at room temperature and filtered using a 0.22μm filter. The phage lysate was then purified using either the PuroMag− Genomic DNA Purification Kit following manufacturer’s instructions under the whole mammalian blood protocol or the Sera-Xtracta Virus/Pathogen Kit according to the manufacturer’s instructions.

Whole genome sequencing was performed using the Nanopore Rapid Sequencing protocol according to the manufacturer’s protocol with minor modifications. Briefly, following DNA extraction of bacteriophages, 10μL DNA was mixed with 1μL of barcoding transposase enzyme (RBK #1-24) and incubated for 2 minutes at 30°C followed by 2 minutes at 80°C and subsequently pooled. Cleanup was performed using a 1X ratio of AMPure XP Beads to sample volume, which was incubated at room temperature for 10 minutes. The DNA was washed with 1mL of 80% ethanol twice and eluted in 20μL of elution buffer (EB for 10 minutes at room temperature. The eluted library was mixed with rapid adaptor (RA) and stored at 4°C until sequencing. Sequencing was performed on a ONT PromethION P2 flow cell nearing end-of-life as defined by the manufacturer (100-1000 pores).

### Basecalling and Assembly

Basecalling was performed using dorado v1.0.1 using super accurate basecalling v5.2 with barcode detection and trimming enabled using either the SQK-RBK114-24 or SQK-RPB114-24 kit name argument for whole genome or direct-from-plaque sequencing respectively. Genomes were assembled using Flye (Kolmogorov et al., 2019) with the ‘--meta’ argument and compared to their reference genomes using FastANI (Jain et al., 2018). Contigs produced by Flye were extracted using Samtools (Danecek et al., 2021).

## Results

### ONT Transposases Retain Activity in Phage Lysate

In order to assess the potential for sequencing from a single plaque, we first determined if the transposase enzymes used in the Rapid Barcoding Kit V14 (SQK-RBK114-24) would be sufficiently active in small volumes of phage lysates isolated from single plaques. Doing so would allow for the direct sequencing of phages from the moment the first plaques appeared without the need for phage amplification and DNA extraction. We theorized that either incompletely packaged phages or damaged capsids would result in free-floating phage DNA or would allow the transposase to access the packaged DNA. Phage plaques were resuspended in 10uL of phage buffer, and sequencing libraries were prepared according to the Rapid sequencing DNA V14 barcoding protocol (Oxford Nanopore Technologies) following the manufacturer’s instructions, substituting purified DNA with a phage suspension.

As shown in Supplementary Figure S2, the use of this library prepared directly from lysate resulted in almost immediate flow cell depletion demonstrated by the disappearance of any sequencing pore (light green) within minutes compared to the expected life span ranging up to 24 hours and the predominant lack of available sequencing pores (light blue). This sequencing attempt resulted in only 735 reads with an average length of 278 bp (PRJNA1338505). To ensure the pore loss was attributable to the library and not a manufacturing or handling defect, we prepared multiple libraries and used flow cells from different batches with similar results. However, while the flow cell produced almost no output, 89 of the 735 short reads matched Arthrobacter phage genomes as their best BLAST hit. This result was promising as the phages were cultivated on an *Arthrobacter globiformis* host, indicating that some phage genetic material was successfully fragmented by the transposase enzyme.

### Amplification of Bacteriophage DNA Using In-House Primers and Different Transposase Sources

As the Rapid PCR Barcoding kit (SQK-RPB114-24) also appears to rely on a transposase inserting the same recognition sequence (GTT TTC GCA TTT ATC GTG AAA CGC TTT CGC GTT TTT CGT GCG CCG CTT CA) into the target DNA, followed by PCR amplification targeted to the inserted sites, we investigated the possibility of using a similar approach with the SQK-

### RBK114-24 provided enzymes

To determine the feasibility of amplifying DNA directly from a plaque and provide a cost-effective prescreen for sufficient phage DNA in an isolation, unmodified, low-cost primers (MuA3’F, Integrated DNA Technologies) were tested as an alternative to the expensive and low-volume of RLB primers provided in the Nanopore Rapid PCR Barcoding Kit which contain the required click-chemistry for sequencing on ONT platforms. This alternative primer (MuA3’F) was designed to bind to the MuA recognition sequence inserted into the DNA by the transposase in the fragmentation step (Figure 1B) (Oxford Nanopore, 2024; Yanagihara & Mizuuchi, 2003). To validate the use of MuA3’F primer instead of RLB primers, 4ng of purified phage DNA was fragmented using the FRM enzyme provided by the Rapid PCR Barcoding Kit. In addition, a negative control with nuclease-free water was incubated with FRM. These fragmented samples were then amplified in the presence of 1εL 10εM MuA3’F or 10εM RLB primers, or a final concentration of 0.5εM primer in a 20εL reaction.

As shown in Figure 2A, both the RLB as well as MuA3’F primers produced an expected smear pattern resulting from amplification of randomly fragmented template DNA. Conversely, the fragmented nuclease-free water with RLB primers and MuA3’F primers nor the fragmented template DNA prior to amplification showed any smearing patterns. Based on the successful amplification of the samples, the MuA3’F primer can potentially serve as a pre-screening, diagnostic or optimization tool. While this primer does not allow for downstream sequencing, it can be used to detect DNA in a plaque isolation as a low-cost alternative to RLB primers.

**Figure 2.**
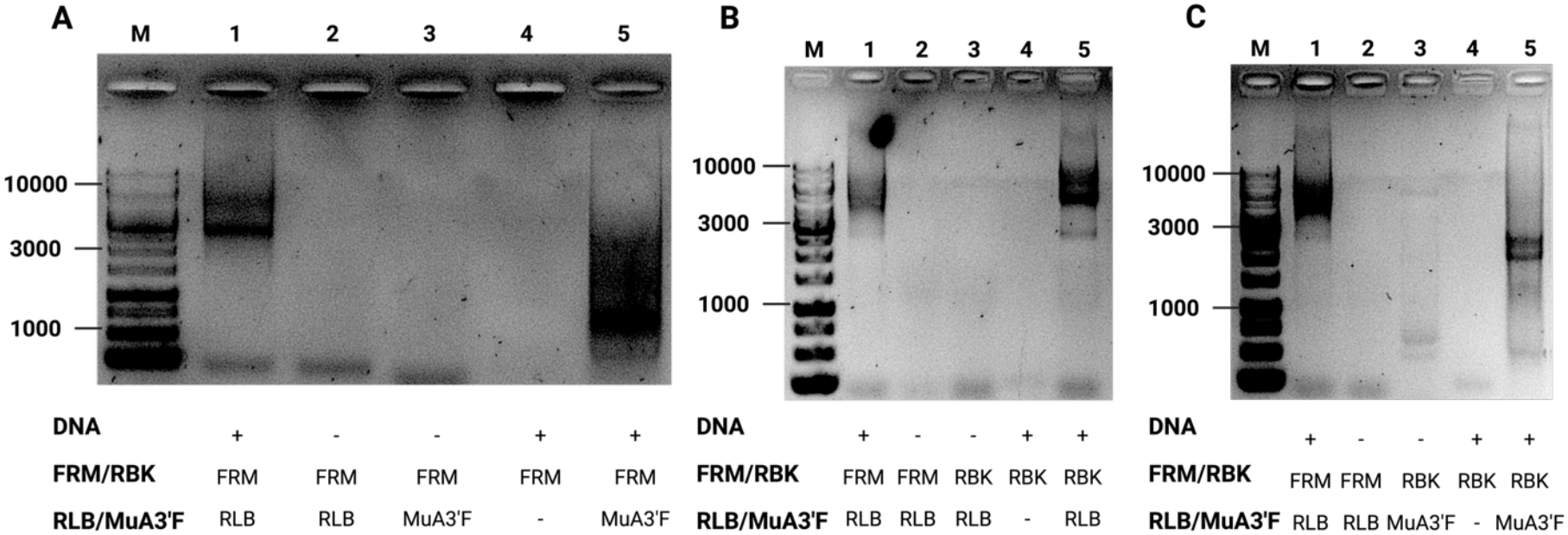
Validation of RBK transposase and MuA3’F primer as functional alternatives for FRM transposase and RLB primers. To establish a cost-effective pre-screening method for direct-from-plaque sequencing, we validated whether the RBK transposase (from the Rapid Barcoding Kit, SQK-RBK114-24) and in-house MuA3’F primers could substitute for the more expensive FRM transposase and RLB primers (from the Rapid PCR Barcoding Kit, SQK-RPB114-24) during optimization steps. Purified phage DNA (4 ng) was fragmented using either transposase, followed by PCR amplification with the indicated primers. All samples were analyzed by agarose gel electrophoresis. **A)**. Purified phage DNA was fragmented using FRM transposase, then amplified with either RLB or MuA3’F primers to compare amplification efficiency. Both RLB (lane 1) and MuA3’F primers (lane 5) produced characteristic smear patterns indicating successful amplification of randomly fragmented DNA. Negative controls (lanes 2, 3) and unamplified template (lane 4) showed no amplification, confirming primer specificity. **B)**. To determine whether the RBK transposase from the Rapid Barcoding Kit could substitute for FRM during optimization, purified phage DNA was fragmented with either transposase and amplified with RLB primers. Both FRM (lane 1) and RBK (lane 5) transposases produced similar amplification smears when combined with RLB primers, demonstrating functional equivalence. Negative controls (lanes 2, 3) showed no amplification. **C.)** To validate whether the cost-effective RBK/MuA3’F combination could predict the outcome of expensive FRM/RLB reactions during optimization, purified phage DNA was processed with both enzyme-primer combinations. The RBK/MuA3’F combination (lane 5) produced amplification patterns comparable to FRM/MuA3’F (lane 1), though with slightly different average fragment sizes.

Additionally, the transposase activity of the RBK enzyme provided by the Rapid Barcoding Kit was compared to the activity of the FRM enzyme provided by the Rapid PCR Barcoding Kit. To determine if the RBK enzyme from the Nanopore Rapid Barcoding Kit serve as a functional alternative to the FRM enzyme from the Nanopore Rapid PCR Barcoding kit, 4ng of purified phage DNA was incubated with 1μL of either FRM or RBK for the fragmentation step of the Nanopore Rapid PCR Barcoding protocol. A negative control using nuclease-free water in place of DNA was additionally incubated with both FRM and RBK. As shown in Figure 2B, gel electrophoresis of the PCR products revealed a smear for the samples amplified using either RBK or FRM with RLB primers, indicating the enzymes can be used interchangeably.

Lastly, we investigated the use of both RBK and MuA3’F replacement within the same sample. Comparable results to the amplification patterns observed when using the Nanopore Rapid PCR Barcoding Kit would allow for prescreening and prediction of PCR outcome before using expensive reagents.

The amplification pattern was tested by incubating 4ng of purified phage DNA with either FRM or RBK, which subsequently underwent amplification in the presence of RLB or MuA3’F primers respectively. As shown in Figure 2C, both the DNA amplified with the FRM/RLB and RBK/MuA3’F combinations showed the expected smears, albeit at slightly different average sizes. Despite this difference in amplified DNA lengths, the combination of RBK and MuA3’F provided comparable results for the presence of amplification. From these results, we determined that processing DNA using the RBK transposase and MuA3’F primer may serve as a suitable method for predicting outcomes of amplification with the Nanopore Rapid PCR Barcoding Kit.

### Reduction of Agar Contaminants Using TE Dilutions

While amplification of purified DNA fragmented by either transposase was possible, and successful fragmentation of phage DNA in lysate was observed directly in a droplet of phage buffer (although at low efficiency, Supplementary Figure S2), we were unable to amplify this fragmented DNA directly. Indeed, the use of the fragmented phage lysate, which previously had a negative effect on flow cell life, also appeared to inhibit the PCR reaction.

As agar contamination is a known PCR inhibitor (Gibb & Wong, 1998), we investigated whether dilution of the plaque pick would increase PCR success rates by diluting potential inhibitors. At the same time, we moved from using phage buffer to 10mM Tris, 0.1 mM EDTA buffer which was also used for successful RBK/MuA3’F amplification using purified DNA to prevent any potential PCR inhibition by phage buffer components. To control for variations in each plaque and phage intrinsic characteristics, a single plaque was isolated from four different bacteriophages (#4, #11, #17, and #18, Supplementary Table S2) and resuspended in 12.5, 25 or 50μL sterile TE buffer. As shown in Supplementary Figure S3A, while partial amplification was seen at the lower dilutions, the 50μL dilution yielded the higher observed amplification. We investigated the dilutions further by comparing amplification of 13 different phages at both 50 and 100μL dilutions. As shown in Supplementary Figure S3B, while the amplification using a 50μL dilution resulted in successful amplification of 10/13 samples, only 3/13 samples had notable amplification using the 100μL dilution, possibly due to over dilution of the template DNA. Due to the increased amplification rate observed with the 50μL dilution, it was determined that 50μL is a suitable volume for plaque resuspension that maintains sufficient DNA concentration while minimizing agar contaminant concentration.

### Prescreening of successful amplification

Following optimization of the TE resuspension volume, and confirmation that MuA3’F primer could amplify transposase fragmented purified DNA (Figure 2B), we investigated the use of MuA3’F as a prescreening tool for the success of barcoded amplification from plaque picks using sequencing (RLB) primers. This prescreen method would allow for a cost-efficient method to determine if the isolated plaque DNA is amenable to PCR amplification and sequencing. To test this, plaques were picked from all phage plates in our student-led collection (Supplementary Table S1) and fragmented using FRM transposase. The fragmented sample was then divided, with 1μL of the sample amplified with MuA3’F and 1μL amplified using RLB barcoded primers.

As shown in Figure 3A, using MuA3’F successfully amplified all samples except #1 and #4. Similarly, all samples amplified with RLB except #1 and #4 showed amplification. While weaker amplification was observed in samples #14 and #21 with RLB as compared to MuA3’F, the same overall pattern of amplification success and failure was observed for both primers used.

**Figure 3.**
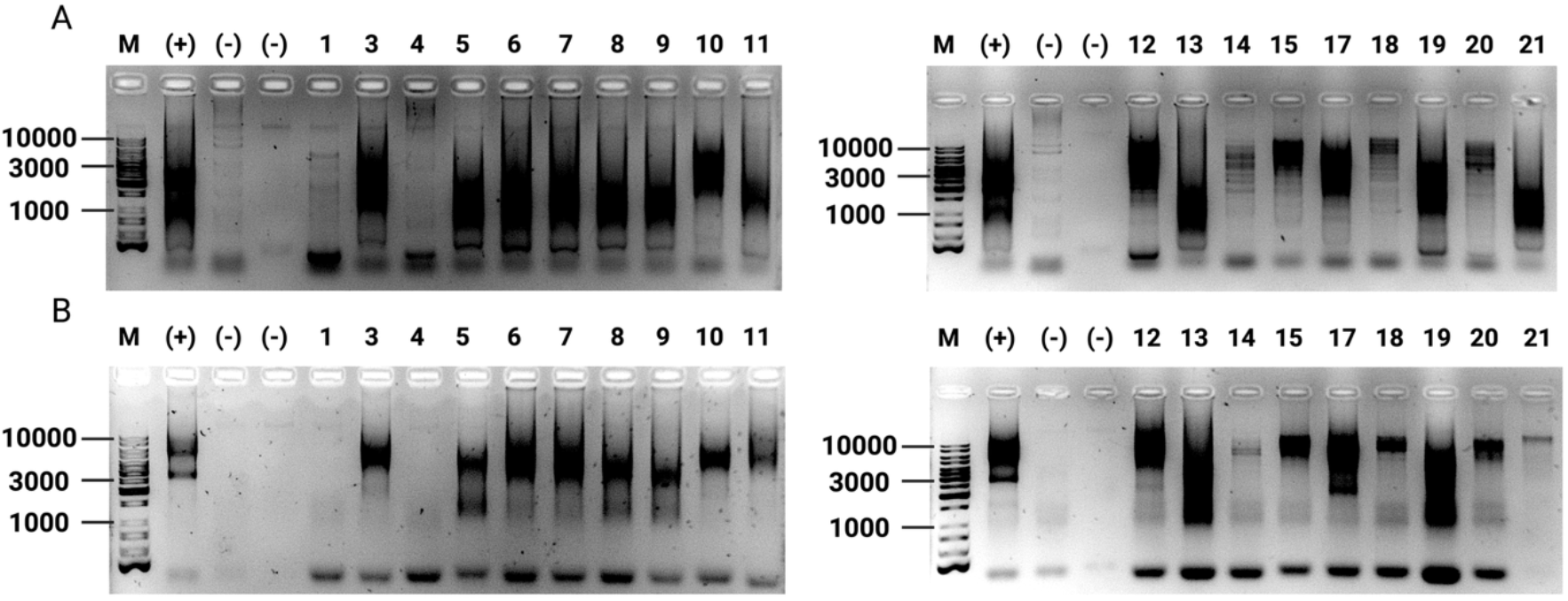
Parallel PCR Amplification Using MuA3’F and RLB Primers. Parallel amplification was performed to determine if PCR using MuA3’F primers could serve as an accurate pre-screen for amplification with sequencing primers. Isolated and resuspended plaques were fragmented using FRM. The fragmented samples were then amplified with either MuA3’F primers **(A)** or RLB primers **(B)**. A positive control (4ng purified phage, lane 2) DNA was amplified, in addition to two negative controls: nuclease-free water that underwent fragmentation and PCR (lane 3), and a fragmented but non-amplified phage sample (lane 4). Characteristic smear patterns indicating successful amplification of randomly fragmented DNA appear in the same lanes for both MuA3’F primers (top) or RLB primers (bottom).

### Whole Genome Sequencing as Reference Genomes

Next, we set out to compare the accuracy of our single plaque sequencing method versus traditional DNA extraction and sequencing protocols. However, only 6 phages available to us from our student-led collection (Supplementary Table S1) had matching high-quality manually assembled and curated reference genomes built from Illumina data through the SEA-PHAGES program (in **bold**, Supplementary Table S2). We therefore first performed Whole Genome Sequencing (WGS) on purified DNA extracted using the phages from this collection on our ONT platform to 1) establish a baseline of errors due to differences in sequencing technologies and 2) to provide a reference genome for all phages analyzed in this study to compare our novel method to.

ONT based whole genome sequencing (ONT-WGS) was performed on 21 DNA extracts using the Nanopore SQK-RBK114.24 Rapid Barcoding Kit. To validate the use of ONT-WGS as reference genomes, the assemblies of the 6 Illumina-sequenced phages were compared to their published genomes: CabbageMan, SJReid, Lewando, MaterMagnus, Boog and Liebe (Table 1). Genome assembly resulted in similar genome lengths for 4/6 phages and an exact length match for one phage genome (MaterMagnus). One phage did not produce assemblies in repeated sequencing experiments (Boog). The average nucleotide identities (ANI) of all phages were above 99.9% with 100% ANI for both CabbageMan and MaterMagnus. Based on the high level of assembly similarity, ONT-based WGS was determined to be a suitable method for obtaining a reference genome for previously non-sequenced bacteriophages.

**Table 1.** Comparison of ONT whole genome sequencing compared to Illumina sequenced reference genomes. Bacteriophages with Illumina-sequenced genomes were cultured to generate lysates followed by DNA extraction and barcoding using the Nanopore Rapid Barcoding Kit. Assembly statistics from Flye assembled genomes and comparisons using FastANI are reported. All ANIs were above 99.9%, apart from Boog (#20), as this phage did not successfully sequence or assemble during whole genome sequencing. *: SJReid was originally assembled at 172839 base pairs due to the presence of terminal repeats and subsequently manually updated by SEA-PHAGES staff.

| Phage (#) | Illumina WGS Length (bp) | ONT WGS Length (bp) | ONT WGS Coverage | Average Nucleotide Identity (%) |
| --- | --- | --- | --- | --- |
| CabbageMan (#4) | 15612 | 15616 | 165 | 100 |
| SJReid (#5) | 193007 (172839)* | 171927 | 11 | 99.9965 |
| Lewando (#18) | 55075 | 55075 | 843 | 99.9946 |
| MaterMagnus (#19) | 53069 | 53069 | 254 | 100 |
| Boog (#20) | 54259 | - | - | - |
| Liebe (#21) | 45803 | 45812 | 105 | 99.9076 |

Following the analysis of the above mentioned ONT-WGS samples, the discovery of identical genomes in multiple phage stocks led to 7 samples being discarded from further analyses (in red, Supplementary Table S2).

### Direct from plaque whole genome sequencing

Having obtained phage reference genomes of sufficient quality to allow for comparisons, we then performed single plaque sequencing on the same phages to compare the resulting genome assemblies. Following amplification prescreen using the MuA3’F primer as described above, the remainder of fragmented samples were amplified and barcoded for sequencing using RLB primers, incubated with RA and sequenced. As shown in Figure 4, successful amplification was attained using the RLB sequencing primers with a 100% success rate.

**Figure 4.**
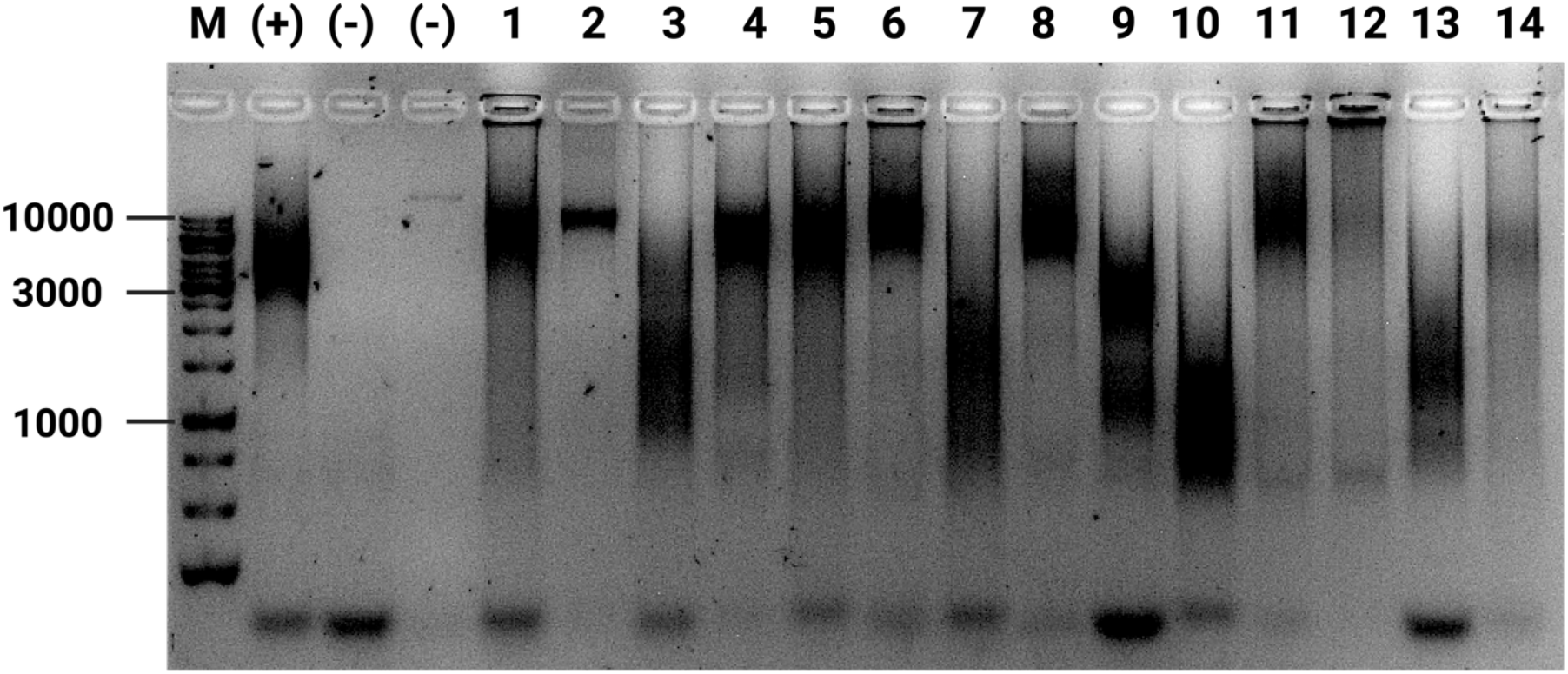
Amplification and barcoding of phage plaque isolates. Isolated and resuspended plaques were fragmented using FRM. The fragmented samples were then amplified using RLB1-14. A positive control (lane 1) with 4ng purified phage DNA and a negative control (lane 2) with nuclease-free water underwent the fragmentation and amplification protocol as well. Phage #1 was also loaded as a pre-amplified fragmented mixture (lane 3). Characteristic smear patterns associated with successful amplification of fragmented DNA appear in all 14 plaque lanes and positive control.

Following sequencing, the single-plaque genomes were assembled and compared to reference genomes originating from traditional DNA extractions and WGS on ONT (Table 2) and Illumina (Table 3). Of the 14 samples sequenced with this method, 7 resulted in genome assemblies with 100% ANI and 7 assemblies with an ANI >99.8% when compared to ONT-WGS results (Table 2). The length of ONT-WGS assemblies as compared to plaque assemblies differed by less than 0.81% for all phages, and averaged a 0.02% difference in length, supporting the feasibility of obtaining highly accurate genome data from a single plaque much more rapidly than traditional methods.

**Table 2.**
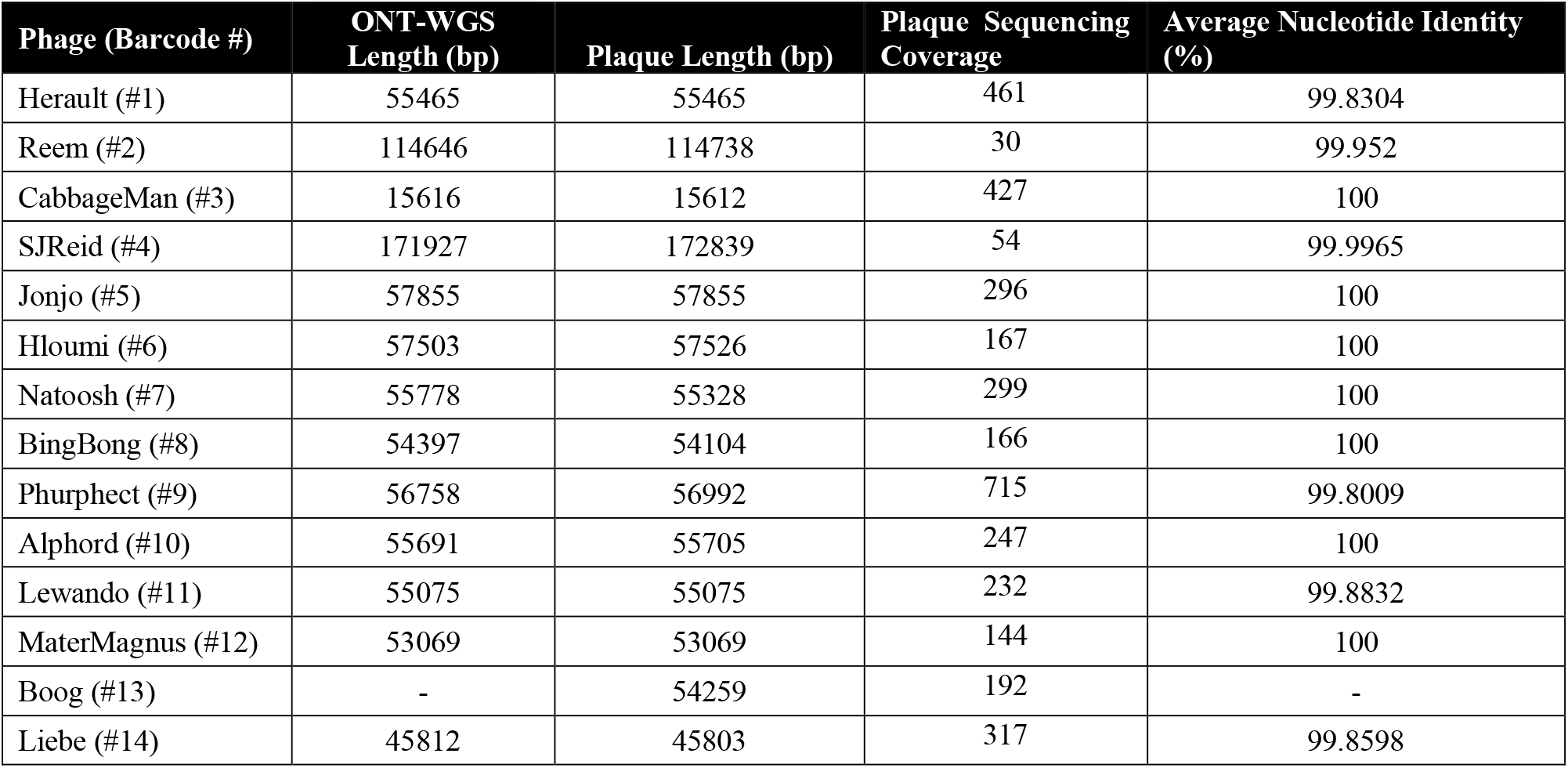
Genome lengths and average nucleotide identities of bacteriophages using ONT whole genome sequencing and direct from plaque sequencing. Sequencing of all phage samples for ONT-WGS was performed as described earlier using purified DNA from phage lysates. Direct from plaque sequencing libraries were prepared as described in Methods. Genomes were assembled using Flye, isolated using samtools, and compared to their ONT-WGS reference genome assembly using FastANI. (As described above, no successful assembly using ONT-WGS was obtained for Boog). Genome lengths are noted in base pairs and average nucleotide identity is represented as a percentage.

**Table 3.** Genome lengths and average nucleotide identities of bacteriophages using direct from plaque sequencing and Illumina sequencing. Direct from plaque sequencing libraries were prepared as described in Methods and compared to their Illumina reference sequence using FastANI. Genome lengths are noted in base pairs and average nucleotide identity is represented as a percentage. **Reported as 99.86% by FastANI, due to a rotation of the circular genome by the Flye assembler. Manual rotation of the last 2100 nucleotides to the start of the genome resulted in a 100% ANI across the entire genome.

| Phage (#) | Illumina Length (bp) | Plaque Length (bp) | Average Nucleotide Identity (%) |
| --- | --- | --- | --- |
| CabbageMan (#3) | 15612 | 15612 | 100 |
| SJReid (#4) | 193007 (172839)* | 172839 | 100 |
| Lewando (#11) | 55075 | 55075 | 100** |
| MaterMagnus (#12) | 53069 | 53069 | 100 |
| Boog (#13) | 54259 | 54259 | 99.9976 |
| Liebe (#14) | 45803 | 45803 | 99.875 |

Importantly however, when compared to Illumina reference genomes, all six phage genomes sequenced through our method matched the exact length of the Illumina reference genome (Table 3). Additionally, 4/6 genomes had an ANI of 100% while one phage had an ANI of 99.9976 % resulting from a single nucleotide change, and one phage at 99.875 % resulting from 3 nucleotide changes (Table 4). Notably, these results seem to indicate that the reference genomes generated from Illumina were more consistent with the single plaque genome assemblies rather than the ONT-WGS assemblies.

**Table 4.** Single nucleotide polymorphisms (SNPs) in plaque assemblies of phages compared to Illumina reference genomes. Genomes were assembled using Flye and compared to their Illumina reference sequence using FastANI. For phages with an ANI below 100%, SNPs were analyzed using mummer. Their location in the reference genome in addition to the nucleotide change, amino acid effect, and gene position.

| Phage (#) | SNP | Location (bp) | Effect (position) | Gene |
| --- | --- | --- | --- | --- |
| Boog (#13) | C → T | 38961 | Glycine (908) → Glycine | 57 |
| Liebe (#14) | G → A | 30523 | Glycine (88) → Arginine | 40 |
| Liebe (#14) | T → C | 42426 | Arginine (15) → Arginine | 62 |
| Liebe (#14) | C → T | 45134 | Alanine (31) → Valine | 68 |

## Discussion

As the pace of phage discovery has until now greatly outpaced the sequencing capabilities, we set out to address this sequencing bottleneck by the development of a technique allowing for the rapid, culture-amplification and DNA extraction-free sequencing of phage genomes from a single plaque as input material. Our study reveals that a simple lysis step exposes enough genetic material to allow transposase-based tagging of the phage genome which then enables sequencing library preparation using PCR primers targeting the transposase inserted region. Although rapid, this method shows identical or nearly identical results to the gold standard of phage amplification, purification and Illumina based sequencing while greatly reducing time and cost requirements.

### Prescreen tool for successful amplification: MuA3’F Primer

An in-house primer designed to bind to the transposase-inserted MuA recognition sequence demonstrated that amplifying phage DNA directly from a plaque is feasible. After fragmentation and amplification of bacteriophage samples with the MuA3’F primer, the smearing observed indicated successful insertion of the MuA recognition sequence into the phage DNA, also confirming the earlier results showing transposase activity towards phage particles (Supplementary Figure S2). This smearing pattern results from the random fragmentation of the genomic DNA by the transposase and is not affected using either FRM or RBK transposase (Figure 2B), allowing the use of left-over RBK enzymes as they are provided in excess of other sequencing reagents (SQK-RBK114-24, ONT). However, as the MuA3’F primer binds to the MuA region inserted by the transposase, amplification of the sample does not result in amplification of the barcode also inserted by the RBK transposase from the Rapid Barcoding Kit, which is situated further upstream in the sequence (Figure 1B). Attempts at generating a primer capable of binding 5’ of the barcode sequence in order to reuse the barcoding from the transposases was unsuccessful. Nevertheless, this inexpensive unbarcoded primer can be used for pre-screening plaque isolations, prior to using expensive barcoded primers from ONT, as the results produced by PCR using MuA3’F closely resembled the results using RLB primers (Figure 2A). The same failures were noted in each PCR run, with all other samples showing smeared bands.

This pre-screening strategy can also be used for further optimization and modification of this protocol, for example, to investigate alternative phage lysis methods. Indeed, the dilution of phage plaques in TE likely leads to osmotic shock in addition to diluting contaminant, increasing the ratio of available DNA versus contaminants. As such, other methods such as freeze-thaw cycling, heating, sonication or other inhibitor removal techniques can possibly replace or complement dilution in TE and allow for successful fragmentation by transposases and amplification.

Additionally, this technique was designed for rapid sequencing and to alleviate the need for the traditional steps needed to extract phage DNA such as generating a large amount of phage lysate. However, as phage lysate is similar in composition to the phage suspensions we obtain from plaque picks, this protocol is also likely to work with existing freezer stocks of phage lysate. A dilution prior to osmotic shock is likely needed as the concentration of phages in amplified phage lysate is much higher than the amount of phage taken up by touching a P20 pipette tip to a plaque.

Lastly, these pre-screening primers may be used to interrogate the application of this method to organisms other than phages. As the MuA transposase appears to remain active in buffers contaminated with agar and bacterial lysate, this method can likely allow for sequencing or any organism susceptible to osmotic shock. The recent publication of a similar protocol by ONT targeted at direct sequencing of *Salmonella* colonies further supports the versatility of this method (Oxford Nanopore, 2025).

### Accuracy of ONT-WGS to Illumina

To create a reference genome for all previously non-sequenced phages, this study found whole genome sequencing using purified phage DNA and the Nanopore Rapid Barcoding Kit to be a suitable alternative to Illumina sequencing. As observed for the five bacteriophages that were successfully sequenced using ONT-WGS, all ANIs exceeded 99.9% (Table 1). Additionally, the genome lengths were identical for three of the phages: CabbageMan, Lewando, and MaterMagnus. SJReid’s genome assembled at 171927 base pairs. While this length differs from the Illumina reference genome, the coverage during sequencing for SJReid was notably low at less than 15. As compared to other samples that assembled with a coverage exceeding 200, the reads were likely insufficient to properly assemble the genome indicating this discrepancy is not due to the accuracy of basecalling in ONT sequencing, but rather a low DNA concentration in the extracted sample. Despite the dissimilar genome length, SJReid’s ONT-WGS ANI as compared to Illumina remained high, at 99.99%. The genome lengths of Liebe and Lewando were not exact matches to their Illumina references, as well. However, they differed by less than 15 base pairs and the ANI remained very high for both. Due to the comparable genome lengths and sequences observed when comparing ONT-WGS to Illumina, it was determined that a suitable reference genome can be produced using ONT-WGS. As native DNA was sequenced for these phages using ONT-WGS, it is possible that phage specific DNA modifications impacted the accuracy of the basecalling as phage genomes are rich in modified bases (Hutinet et al., 2021). Additionally, phage Liebe was first sequenced in 2017 and has undergone several passaging rounds as it is used as a positive control in the SEA-PHAGES program. It is therefore possible this phage accumulated mutations from the original stock that was used to generate the reference Illumina genome.

### Sequencing of single plaques

Through direct isolation of individual plaques, it was observed that amplification, sequencing, and genome assembly were successful for all phage samples through use of the Nanopore Rapid PCR Barcoding Kit. Utilizing a pre-screen of amplification with the MuA3’F primer allowed for the identification of poorly isolated phage DNA and therefore allowed for re-isolation of select phages. Following amplification, sequencing proved the accuracy of PCR-based sequencing as compared to both Illumina and ONT-WGS assemblies. All calculated ANIs exceeded 99.80% for both comparisons to Illumina and ONT-WGS (Table 2, 3). The mean ANI for PCR assemblies using their Illumina reference genomes was 99.98%, with an identical genome length match for all six samples, apart from the terminal repeat for SJReid (which was also originally truncated in its Illumina assembly, prior to manual curation).

Notably, it was observed that the genome lengths assembled using this method had a much higher match to Illumina genome lengths are compared to whole genome sequencing. This increased match is likely due to the introduction of a PCR step, as phages are frequently modified and PCR amplification results in DNA with only the four canonical bases. A limitation of this technique compared to ONT-WGS is therefore the study of DNA modifications, similar to Illumina sequenced genomes.

While WGS of DNA obtained from a single plaque have been previously reported (DePew et al., 2013; Kot et al., 2014), the methods described makes use of several enzymatic steps or overnight incubation and only produce a sequenceable amount of genetic material. This amplified material still must be transformed into sequencing libraries followed by Illumina sequencing while our protocol generates ready to sequence ONT libraries and allows for sequencing on ONT platforms which come with the advantages of live data acquisition, portability and controls during sequencing such as adaptive sampling (discussed below).

### Bacterial contaminants

While the phage genomes were frequently the only genomes assembled from the sample, some leftover bacterial DNA present in the sample can also be amplified by the PCR step. Due to the natural increase in phage copy number prior to bacterial lysis, however, the phage genomes are almost always dominant. Extracting phage genomes from complex assemblies is easily done using samtools post-assembly. Alternatively, as the bacterial hosts are generally known during phage screens, the reads originating from the hosts can be removed either during sequencing, thanks to ONT’s adaptive sampling tool which allows for the physical rejection of reads belonging to certain genomes, or post-sequencing but prior to assembly by mapping reads to a reference host genome (Danecek et al., 2021; Martin et al., 2022). Due to the highly sensitive nature of this technique however, any contaminating DNA will be amplified and sequenced therefore all steps performed prior to tagmentation require good aseptic technique to avoid other bacterial, environmental, or human DNA from contaminating the sample.

### Further uses

The technique of direct isolation is also a quick and effective tool to rapidly screen for contamination. As our initial screening with RLB primers revealed a contaminating phage in select samples (Supplementary Table S2) in all freezer stocks of CabbageMan, we performed sequencing of multiple plaques per stock. Using this technique, we were able to identify pure from mixed stocks and were able to rescue a contaminated sample rapidly and at low cost.

While the RLB primers provided by the SQK-RBK offer a rapid method of sequencing up to 24 phages per flow cell, higher multiplexing is possible due to the small genome size of phages. Hundreds of custom barcodes (Srivathsan et al., 2024) could be added to the MuA3’F sequence to sequence more phages at once to reduce sequencing costs and sequence more pages at once. This higher multiplexing however would come at a slight expense of time as the unique chemistry of RLB primers allows for the rapid attachment to the sequencing adapter immediately post-PCR while custom barcoded primers would require a subsequent enzymatic ligation.

### Concluding remarks

The increased similarity in assembly as well as compatibility with impure samples further supports this sequencing method as an accurate and fast approach to DNA sequencing. To further confirm this, additional sequencing should be performed on other Illumina-sequenced bacteriophages.

The high sequence similarity as well as complete amplification for all samples suggests that this method of DNA isolation and sequencing could be widely applicable to diverse bacteriophages.

Overall, this experiment demonstrates the feasibility of direct plaque isolation and PCR amplification of bacteriophages to produce highly accurate genome sequences that can be used for downstream bioinformatic applications.

## Data availability

Whole-genome sequences have been deposited into GenBank and are publicly available as of the date of publication under the NCBI BioProject: PRJNA1338505

## Acknowledgments

This research was supported by a grant from the Canadian Institute of Health Research (CIHR 195796) to PL. The phage collection and isolation were performed by the 2022 and 2023 cohorts of SEA-PHAGES students (Supplementary Table S1) at McGill University under the supervision of Michael Shamash and Dr Corinne Maurice.

## References

Danecek, P., Bonfield, J. K., Liddle, J., Marshall, J., Ohan, V., Pollard, M. O., Whitwham, A., Keane, T., McCarthy, S. A., Davies, R. M., & Li, H. (2021). Twelve years of SAMtools and BCFtools. Gigascience, 10(2). 10.1093/gigascience/giab008

DePew, J., Zhou, B., McCorrison, J. M., Wentworth, D. E., Purushe, J., Koroleva, G., & Fouts, D. E. (2013). Sequencing viral genomes from a single isolated plaque. Virology Journal, 10(1), 181. 10.1186/1743-422X-10-181

Elfadadny, A., Ragab, R. F., Abou Shehata, M. A., Elfadadny, M. R., Farag, A., Abd El-Aziz, A.H., & Khalifa, H. O. (2024). Exploring Bacteriophage Applications in Medicine and Beyond. Acta Microbiologica Hellenica, 69(3), 167–179. https://www.mdpi.com/2813-9054/69/3/16

Gauthier, C. H., & Hatfull, G. F. (2024). A Bioinformatic Ecosystem for Bacteriophage Genomics: PhaMMSeqs, Phamerator, pdm_utils, PhagesDB, DEPhT, and PhamClust. Viruses, 16(8). 10.3390/v16081278

Gibb, A. P., & Wong, S. (1998). Inhibition of PCR by agar from bacteriological transport media. J Clin Microbiol, 36(1), 275–276. 10.1128/jcm.36.1.275-276.1998

González-Escalona, N., Allard, M. A., Brown, E. W., Sharma, S., & Hoffmann, M. (2019). Nanopore sequencing for fast determination of plasmids, phages, virulence markers, and antimicrobial resistance genes in Shiga toxin-producing Escherichia coli. PLoS One, 14(7), e0220494. 10.1371/journal.pone.0220494

Heller, D. M., Sivanathan, V., Asai, D. J., & Hatfull, G. F. (2024). SEA-PHAGES and SEAGENES: Advancing Virology and Science Education. Annu Rev Virol, 11(1), 1–20. 10.1146/annurev-virology-113023-110757

Hutinet, G., Lee, Y. J., de Crécy-Lagard, V., & Weigele, P. R. (2021). Hypermodified DNA in Viruses of E. coli and Salmonella. EcoSal Plus, 9(2), eESP00282019. 10.1128/ecosalplus.ESP-0028-2019

Jain, C., Rodriguez, R. L., Phillippy, A. M., Konstantinidis, K. T., & Aluru, S. (2018). High throughput ANI analysis of 90K prokaryotic genomes reveals clear species boundaries. Nat Commun, 9(1), 5114. 10.1038/s41467-018-07641-9

Kolmogorov, M., Yuan, J., Lin, Y., & Pevzner, P. A. (2019). Assembly of long, error-prone reads using repeat graphs. Nat Biotechnol, 37(5), 540–546. 10.1038/s41587-019-0072-8

Kot, W., Vogensen, F. K., Sørensen, S. J., & Hansen, L. H. (2014). DPS - a rapid method for genome sequencing of DNA-containing bacteriophages directly from a single plaque. J Virol Methods, 196, 152–156. 10.1016/j.jviromet.2013.10.040

Martin, S., Heavens, D., Lan, Y., Horsfield, S., Clark, M. D., & Leggett, R. M. (2022). Nanopore adaptive sampling: a tool for enrichment of low abundance species in metagenomic samples. Genome Biol, 23(1), 11. 10.1186/s13059-021-02582-x

Mushegian, A. R. (2020). Are There 10(31) Virus Particles on Earth, or More, or Fewer? J Bacteriol, 202(9). 10.1128/jb.00052-20

Oxford Nanopore, T. (2024). Chemistry Technical Document. https://nanoporetech.com/document/chemistry-technical-document

Oxford Nanopore, T. (2025). Direct-from-colony rapid microbial sequencing for Salmonella serotyping using SQK-RPB114.24. https://nanoporetech.com/document/direct-from-colony-microbial-sequencing-rapid-salmonella-rpb114-24

Pottie, I., Vázquez Fernández, R., Van de Wiele, T., & Briers, Y. (2024). Phage lysins for intestinal microbiome modulation: current challenges and enabling techniques. Gut Microbes, 16(1), 2387144. 10.1080/19490976.2024.2387144

Rihtman, B., Meaden, S., Clokie, M. R., Koskella, B., & Millard, A. D. (2016). Assessing Illumina technology for the high-throughput sequencing of bacteriophage genomes. PeerJ, 4, e2055. 10.7717/peerj.2055

Salam, M. A., Al-Amin, M. Y., Salam, M. T., Pawar, J. S., Akhter, N., Rabaan, A. A., & Alqumber, M. A. A. (2023). Antimicrobial Resistance: A Growing Serious Threat for Global Public Health. Healthcare (Basel), 11(13). 10.3390/healthcare11131946

Srivathsan, A., Feng, V., Suárez, D., Emerson, B., & Meier, R. (2024). ONTbarcoder 2.0: rapid species discovery and identification with real-time barcoding facilitated by Oxford Nanopore R10.4. Cladistics, 40(2), 192–203. 10.1111/cla.12566

Yanagihara, K., & Mizuuchi, K. (2003). Progressive structural transitions within Mu transpositional complexes. Mol Cell, 11(1), 215–224. 10.1016/s1097-2765(02)00796-7

Zhang, M., Zhang, T., Yu, M., Chen, Y. L., & Jin, M. (2022). The Life Cycle Transitions of Temperate Phages: Regulating Factors and Potential Ecological Implications. Viruses, 14(9). 10.3390/v14091904

Zhang, T., Li, H., Jiang, M., Hou, H., Gao, Y., Li, Y., Wang, F., Wang, J., Peng, K., & Liu, Y. X. (2024). Nanopore sequencing: flourishing in its teenage years. J Genet Genomics, 51(12), 1361–1374. 10.1016/j.jgg.2024.09.007

Zheng, P., Zhou, C., Ding, Y., Liu, B., Lu, L., Zhu, F., & Duan, S. (2023). Nanopore sequencing technology and its applications. MedComm (2020), 4(4), e316. 10.1002/mco2.316

