## Supplementary Information for "DNA extraction free whole genome sequencing of bacteriophage genomes from a single plaque"

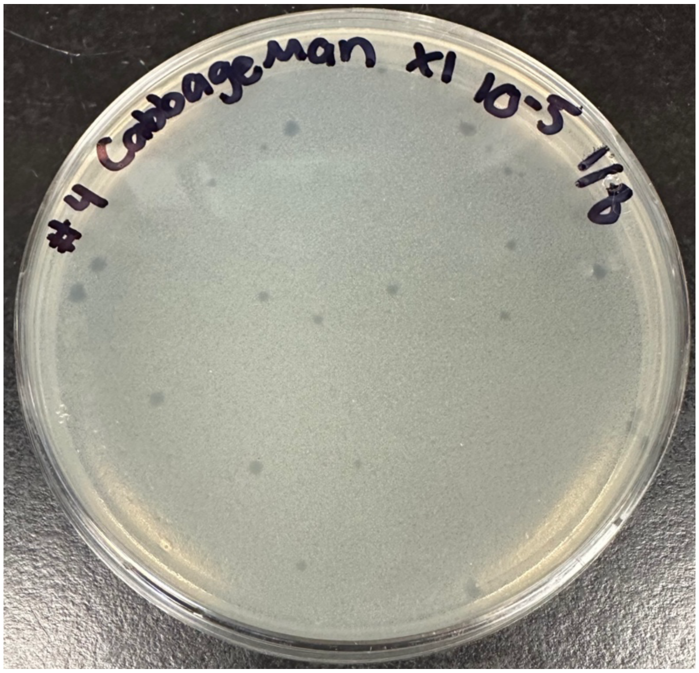


**Supplementary Figure S1. Plaque Assay Example**. Plaque assays were made for all phages used for the rapid PCR barcoding protocol. To produce plates with individual plaques, serially diluted phage stocks were incubated with 100-200µL of the host bacteria *A. globiformis*. Incubation at 30°C for 24-48 hours resulted in bacterial lawns with isolated phage plaques. The figure shows a plaque assay produced using the phage CabbageMan, with plaques resembling small clear circles. Individual plaques were picked and resuspended for use in the rapid PCR barcoding protocol.

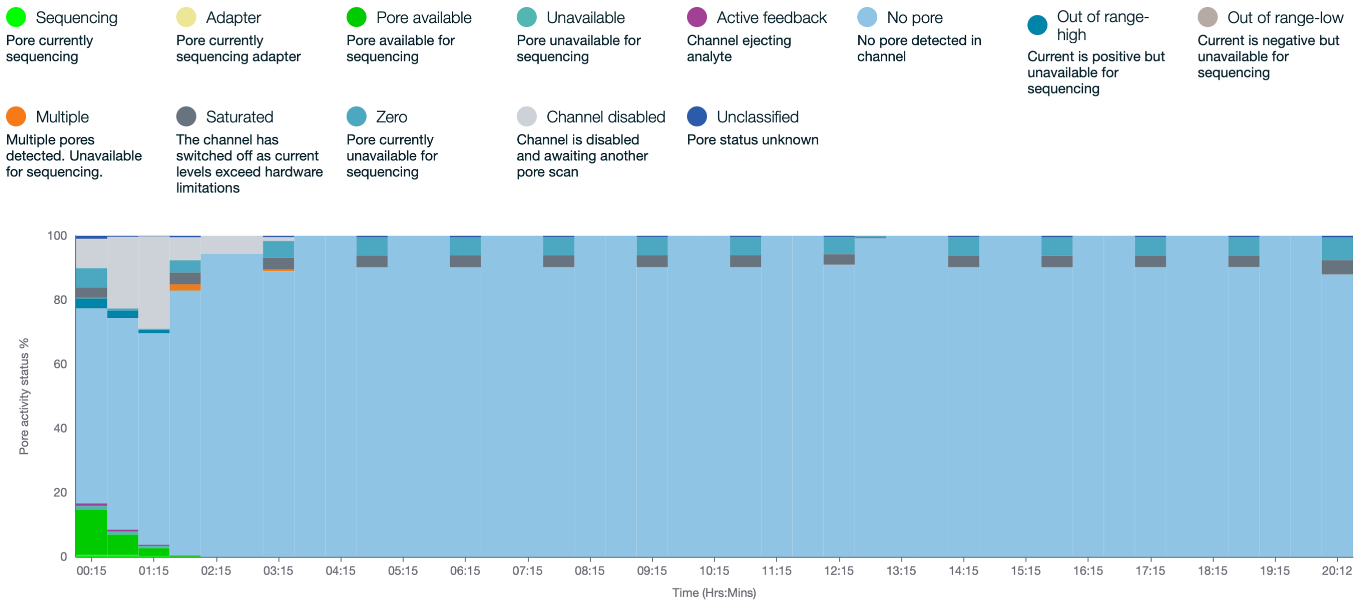


**Supplementary Figure S2. Pore activity report during sequencing from phage plaque suspensions.**

Phage plaques were resuspended in Phage Buffer and incubated with 1uL of RBK transposase enzyme from SQK-RBK114-96 at 30°C for 2 minutes followed by 80°C for 2 minutes. Libraries were pooled and loaded according to manufacturer's instructions on a Flongle device. The rated life span of the flow cell is 24h. Activity report generated using MinKNOW v24.02.8.

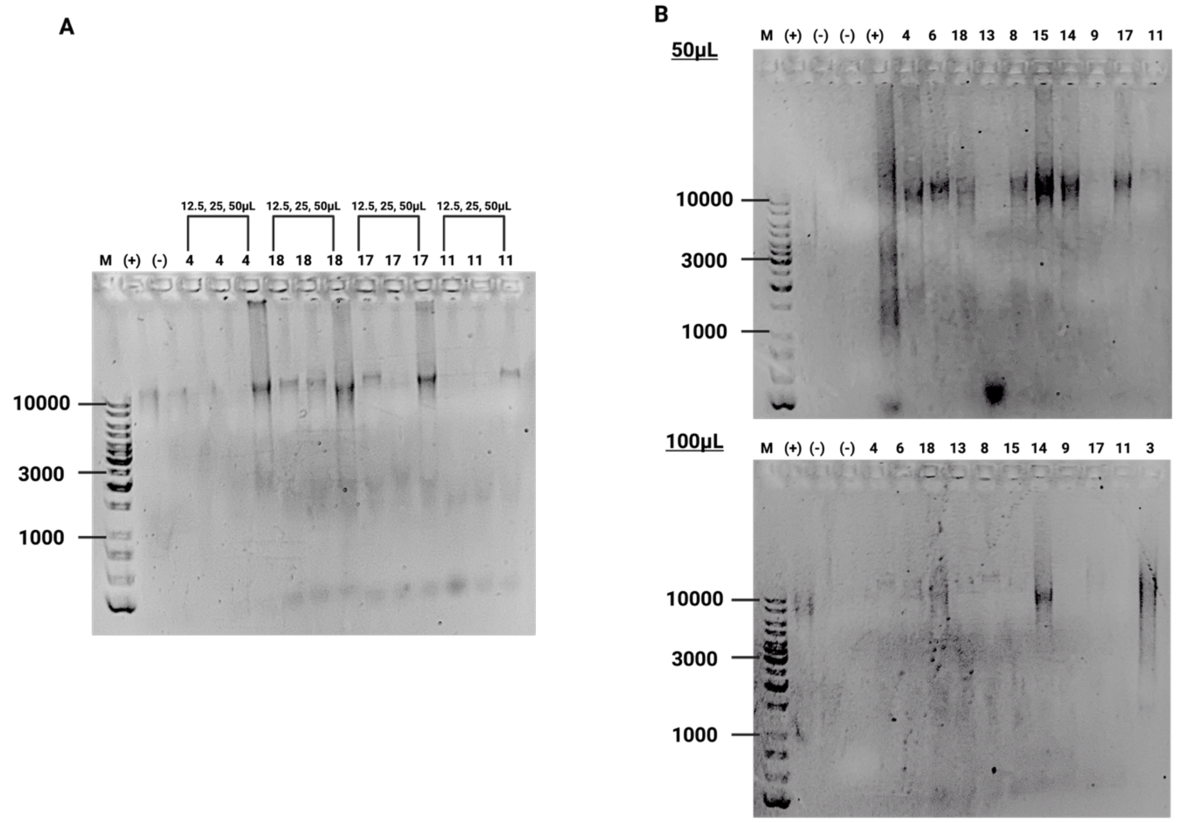


**Supplementary Figure S3. Optimization of TE Resuspension for Plaque Isolation. A)** Individual plaques from phages (#4, 11, 17, and 18) were isolated and resuspended in 12.5, 25 or 50µLof TE buffer and amplified. B) Individual plaques from phages (#3, 4, 6, 8, 9, 11, 13, 14, 15, 17, and 18) were isolated and resuspended in 50 (top) or 100µL (bottom) of TE buffer prior to amplification. A positive control (4ng purified phage DNA) and a negative control (nuclease-free water) were additionally included. An additional positive control (4ng purified phage DNA) was added with an increased final primer concentration (0.5µM), as opposed to all other samples which had a final RLB primer concentration of 0.1µM.

**Supplementary Table S1. Phage Samples used in this study**

| Phage | Isolated by |
|---|---|
| Herault | Vincent Duquette |
| DuSol | Daniel Rownaghi |
| Reem | Reem Araji |
| CabbageMan | Ximena Iraheta |
| SJReid | Hannah Kumar |
| Jonjo | Rishabh Johri |
| Hloumi | Taline Bawab |

| | |
|---|---|
| Natoosh | Natalie Abuelsamen |
| Ratto | Liam Stanley |
| Darami | Victoria Woo |
| Rosales | Brenna Fox |
| BingBong | Maiya Hernandez-Morrison |
| Zinzli | Andrew Dayton |
| Phurrphect | Charlotte Zickmantel |
| Zorgle | Maya Zandstra |
| FINN1505 | Carolyn Bawden |
| Alphord | Jackie Alphord |
| Lewando | Maximilian Crosby |
| MaterMagnus | Chloe Nyiligira, Norah Driscoll |
| Boog | Paige Andrusko, Lynn Gedeon |
| Liebe | Audrey Jonas |

**Supplementary Table S2. Data Sets 1 and 2 for Phages**

| Phage | Barcode Number Data Set 1 | Barcode Number Data Set 2 |
|---|---|---|
| Herault | 1 | 1 |
| DuSol | 2 | - |
| Reem | 3 | 2 |
| **CabbageMan** | 4 | 3 |
| **SJReid** | 5 | 4 |
| Jonjo | 6 | 5 |
| Hloumi | 7 | 6 |
| Natoosh | 8 | 7 |
| Ratto | 9 | - |
| Darami | 10 | - |
| Rosales | 11 | - |
| BingBong | 12 | 8 |
| Zinzli | 13 | - |
| Phurrphect | 14 | 9 |
| Zorgle | 15 | - |
| FINN1505 | 16 | - |
| Alphord | 17 | 10 |
| **Lewando** | 18 | 11 |

| | | |
|---|---|---|
| **MaterMagnus** | 19 | 12 |
| **Boog** | 20 | 13 |
| **Liebe** | 21 | 14 |

*Phages in red all assembled as the same genome in ONT-WGS, and were removed from the second data set as duplicates or contaminated stocks. Phages in blue were not successfully cultured for either webbed plates or plaque assays and were therefore removed from the second data set as well. Phages in **bold** have been previously sequenced using Illumina platforms as part of the SEA-PHAGES program and have genomic data available online.
